# Candidate effector proteins from the maize tar spot pathogen *Phyllachora maydis* localize to diverse plant cell compartments

**DOI:** 10.1101/2022.05.24.492667

**Authors:** Matthew Helm, Raksha Singh, Rachel Hiles, Namrata Jaiswal, Ariana Myers, Anjali S. Iyer-Pascuzzi, Stephen B. Goodwin

**Affiliations:** Crop Production and Pest Control Research Unit, U.S. Department of Agriculture-Agricultural Research Service (USDA-ARS), West Lafayette, IN 47907, U.S.A; Department of Botany and Plant Pathology, Purdue University, West Lafayette, IN 47907, U.S.A

**Keywords:** *Phyllachora maydis*, tar spot, effectors, localization, plasma membrane, nucleus, nucleolus, chloroplasts

## Abstract

Most fungal pathogens secrete effector proteins into host cells to modulate their immune responses, thereby promoting pathogenesis and fungal growth. One such fungal pathogen is the ascomycete *Phyllachora maydis*, which causes tar spot disease on leaves of maize (*Zea mays*). Sequencing of the *P. maydis* genome revealed 462 putatively secreted proteins of which 40 contain expected effector-like sequence characteristics. However, the subcellular compartments targeted by *P. maydis* effector candidate (PmECs) proteins remain unknown and it will be important to prioritize them for further functional characterization. To test the hypothesis that PmECs target diverse subcellular compartments, cellular locations of super Yellow Fluorescent Protein (sYFP)-tagged *P. maydis* effector candidate proteins were identified using a *Nicotiana benthamiana*-based heterologous expression system. Immunoblot analyses showed that most of the PmEC-fluorescent protein fusions accumulated protein in *N. benthamiana*, indicating the candidate effectors could be expressed in dicot leaf cells. Laser-scanning confocal microscopy of *N. benthamiana* epidermal cells revealed most of the *P. maydis* putative effectors localized to the nucleus and cytosol. One candidate effector, PmEC01597, localized to multiple subcellular compartments including the nucleus, nucleolus, and plasma membrane while an additional putative effector, PmEC03792, preferentially labelled both the nucleus and nucleolus. Intriguingly, one candidate effector, PmEC04573, consistently localized to the stroma of chloroplasts as well as stroma-containing tubules (stromules). Collectively, these data suggest effector candidate proteins from *P. maydis* target diverse cellular organelles and may thus provide valuable insights into their putative functions as well as host processes potentially manipulated by this fungal pathogen.

## INTRODUCTION

Plant pathogens secrete virulence proteins known as effectors to modulate host immune responses that often have a functional role in facilitating infection (Jones and Dangl, 2006; Kamoun, 2007; Wang et al., 2011; Zipfel, 2014). Secreted effectors can be either retained in the plant extracellular space (apoplastic effectors) or translocated into host cells (cytoplasmic effectors) and can localize to diverse subcellular compartments (Lorrain et al., 2018; Whisson et al., 2016). For example, an effector protein from the oomycete *Phytophthora infestans*, PITG_04097, localizes to the host nucleus, and such nuclear localization is required for suppression of host defense responses and pathogen virulence (Zheng et al., 2014). The *Pseudomonas syringae* effector HopG1, which targets the mitochondria, suppresses host defense responses and promotes cell death in Arabidopsis (Rodriguez-Puerto et al., 2022). The *Magnaporthe oryzae* effector, AVR-Pii, is localized to the host cytosol where it suppresses host production of reactive oxygen species through its inhibition of the rice NADP-malic enzyme2, thereby disrupting immunity to this fungal pathogen (Singh et al., 2016). Elucidating how crop pathogen effectors function in host cells is critical, in part, for understanding pathogenicity and virulence mechanisms of fungal pathogens for which control strategies are currently limited.

*Phyllachora maydis* is a foliar, ascomycete fungal pathogen that causes tar spot disease on maize (*Zea mays* subsp. *mays*) (Rocco da Silva et al., 2021; Ruhl et al., 2016; Valle-Torres et al., 2020). Though endemic to Central and South America, *P. maydis* was recently identified in the continental United States in 2015 and has since spread to most maize production regions, indicating this fungal pathogen is capable of significant global expansion (Mottaleb et al., 2019; Ruhl et al., 2016; Valle-Torres et al., 2020). Notably, *P. maydis* has been shown to significantly reduce maize yields especially under favorable environmental conditions, imposing severe financial constraints to growers (Mueller et al., 2020; Valle-Torres et al., 2020). Temperate-derived maize inbreds and commercial hybrids provide only partial resistance to *P. maydis,* and no fully resistant maize cultivar has been identified (Telenko et al., 2019). For these reasons, *P. maydis* is now considered one of the most economically important foliar pathogens of maize in the U.S. (Mueller et al., 2020; Rocco da Silva et al., 2021; Valle-Torres et al., 2020).

To gain initial insights into *P. maydis* virulence mechanisms, Telenko and colleagues provided its first draft genome sequence (Telenko et al., 2020). Analysis of the *P. maydis* genome revealed 462 proteins comprising the predicted secretome, of which 59 contain effector-like sequence characteristics as predicted by EffectorP (v2.0) (Telenko et al., 2020). To date, our understanding of how *P. maydis* utilizes its effector repertoire to promote virulence as well as the subcellular compartments targeted by these putative effectors remains limited even though this fungal pathogen represents a serious economic concern for maize growers (Helm et al., 2022; Mueller et al., 2020; Valle-Torres et al., 2020). The inability to culture or genetically manipulate *P. maydis* (Rocco da Silva et al., 2021; Valle-Torres et al., 2020) substantially hinders investigations aimed at characterizing its effector repertoire (Helm et al., 2022). To circumvent these limitations, the field of effector biology utilizes a surrogate plant system to express epitope-tagged candidate effectors directly inside leaf cells using *Agrobacterium*-mediated infiltration (agroinfiltration) (Lorrain et al., 2018). *Nicotiana benthamiana* is a well established and extensively used model plant for heterologous expression of crop pathogen effectors and has been used to investigate the subcellular compartments targeted by putative effector proteins produced by filamentous fungal pathogens (Alfano, 2009; Figueroa et al., 2021; Lorrain et al., 2018; Ma et al., 2012; Petre et al., 2017; Win et al., 2011; Dinne et al., 2021).

In the present study, we refined previous effector predictions performed by Telenko et al. (2020) using EffectorP (v3.0) as well as additional selection criteria including i) protein size less than 300 amino acids; ii) presence of a signal peptide (as predicted by SignalP v6.0); and iii) lack of a transmembrane domain. We discovered that among the 59 proteins originally identified by Telenko and colleagues (2020), 40 contain effector-like protein characteristics that fulfilled our more selective criteria. Intriguingly, several of the effector candidates from *P. maydis* encode subcellular targeting sequences including nuclear localization signals (NLS) and chloroplast and mitochondrial transit peptides, suggesting they may be targeted to specific subcellular locations. To test this hypothesis, we investigated the subcellular compartments targeted by *P. maydis* effector candidate proteins (PmECs) using a *Nicotiana benthamiana*-based heterologous expression system. Laser-scanning confocal microscopy of *N. benthamiana* epidermal cells revealed most of the *P. maydis* putative effectors localized to the nucleus and cytosol. However, one candidate effector, PmEC01597, consistently localized to multiple subcellular compartments including the nucleus, nucleolus, and plasma membrane while an additional putative effector, PmEC03792, preferentially labelled both the nucleus and nucleolus. Intriguingly, the candidate effector, PmEC04573, consistently localized to the stroma of chloroplasts as well as stroma-containing tubules (stromules). These data indicate effector candidate proteins from *P. maydis* target diverse cellular organelles and thus lay the foundation for future studies to investigate their putative functions as well as host processes potentially manipulated by this fungal pathogen.

## MATERIALS AND METHODS

### Plant growth conditions

*Nicotiana benthamiana* seeds were sown in plastic pots containing either ProMix or Berger Seed and Propagation Mix supplemented with Osmocote slow-release fertilizer (14-14-14). Plants were maintained in a growth chamber with a 16:8 h photoperiod (light:dark) at 24°C with light and 20°C in the dark and 60% humidity with average light intensities at plant height of 120 µmols/m^2^/s.

### *In silico* selection of candidate effectors from *P. maydis* Indiana isolate PM-01

To select candidate effector proteins, we began with the effector predictions generated previously by Telenko et al., (2020) from the predicted *P. maydis* secretome. We extracted all 59 candidate effector protein sequences and used EffectorP (v3.0) (Sperschneider and Dodds, 2022) (http://effectorp.csiro.au/) to further improve the effector prediction performed by Telenko et al. (2020). We also employed SignalP (v6.0) (Teufel et al., 2022) (https://services.healthtech.dtu.dk/service.php?SignalP) to predict the presence of signal peptide sequences. LOCALIZER (v1.0) (Sperschneider et al., 2017) (http://localizer.csiro.au/) was used to predict the subcellular localizations of the putative effectors. TMHMM (v2.0; https://services.healthtech.dtu.dk/service.php?TMHMM-2.0) was used to predict transmembrane helices within the candidate effector proteins. NoD (Scott et al., 2011) (http://www.compbio.dundee.ac.uk/www-nod/) was used to predict the presence of predicted Nucleolar targeting signal (NoLS). The refined catalog of candidate effector proteins as determined by our *in silico* pipeline (Figure 1) is shown in Table 1.

**Figure 1.**
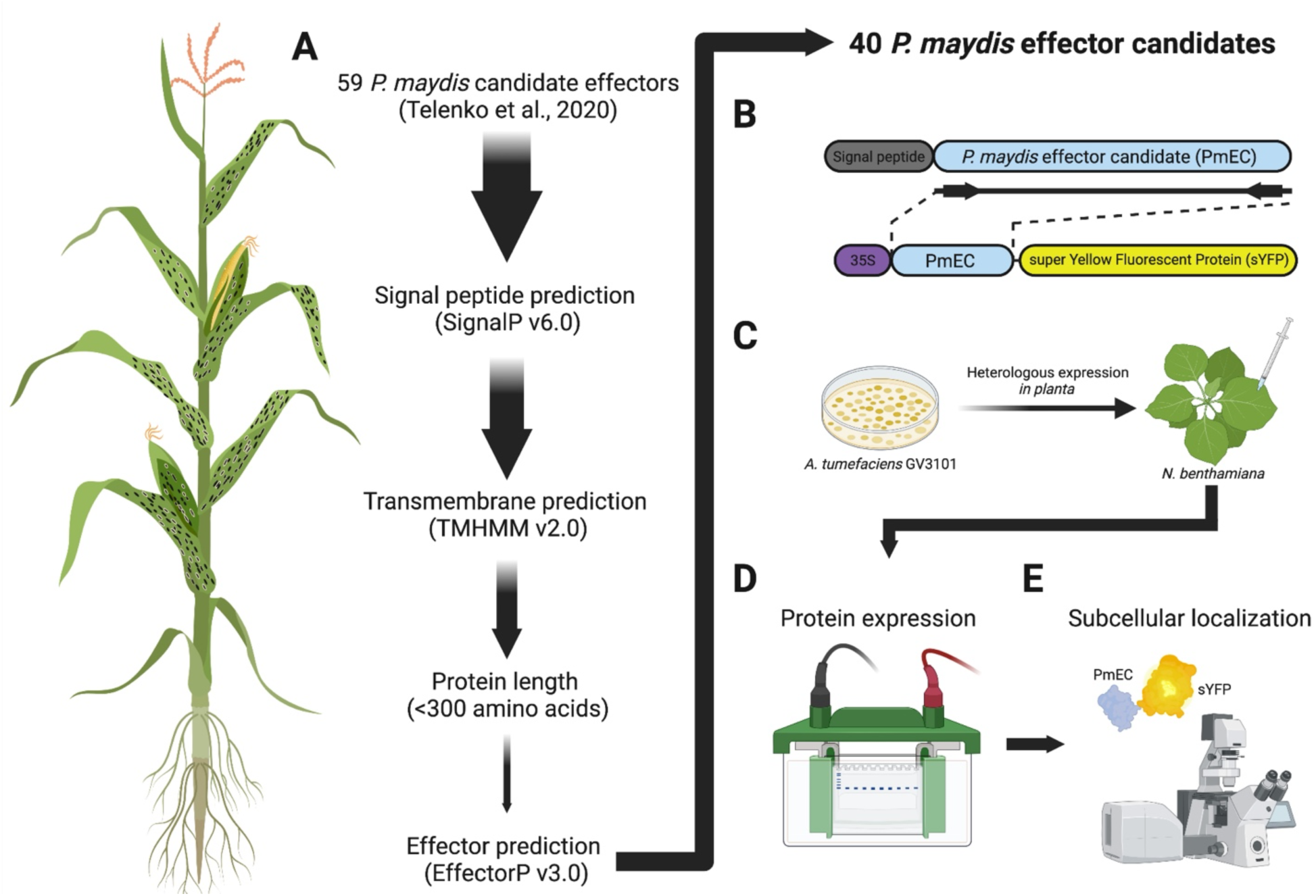
Schematic overview of the selection and subsequent analyses of *Phyllachora maydis* candidate effectors. **A)** The *P. maydis* effector candidates investigated in this study were selected using the aforementioned selection criteria. **B)** The predicted open reading frames (ORFs) of each of the 40 candidate effectors, without their predicted signal peptides, were synthesized and fused to the N terminus of super Yellow Fluorescent Protein (sYFP) and recombined into the plant expression binary vector pEarleyGate100 (pEG100) using a multisite Gateway cloning strategy. **C)** The resulting *P. maydis* effector-fluorescent protein fusion (PmEC:sYFP) constructs were inserted into *Agrobacterium tumefaciens* for subsequent *Nicotiana benthamiana*-based heterologous expression assays. **D)** Immunoblot analyses were used to assess expression of the PmEC-fluorescent protein fusions. **E)** Laser-scanning confocal microscopy was used to assess the live-cell subcellular localization patterns in *N. benthamiana* epidermal cells. Figure created with www.BioRender.com.

**Table 1.** *Phyllachora maydis* effector candidates (PmEC) investigated in this study.

| Protein ID<sup>a</sup> | Amino acids<sup>b</sup> | Signal peptide<sup>c</sup> | Protein size (kDa)<sup>d</sup> | LOCALIZER prediction<sup>e</sup> | *in planta* localization<sup>f</sup> |
| --- | --- | --- | --- | --- | --- |
| PmEC00197 | 262 | 16-17 | 27.3 |  | Nucleus and cytosol |
| PmEC00457 | 156 | 20-21 | 14.9 |  | Nucleus and cytosol |
| PmEC00684 | 140 | 30-31 | 13.3 |  | Nucleus and cytosol |
| PmEC00848 | 177 | 20-21 | 17.2 | Chloroplast (56-78)<br>Mitochondria (67-87) | Nucleus and cytosol |
| PmEC01139 | 201 | 24-25 | 19.3 |  | Nucleus and cytosol |
| PmEC01169 | 55 | 20-21 | 3.9 |  | Nucleus and cytosol |
| PmEC01289 | 109 | 20-21 | 10.2 |  | Nucleus and cytosol |
| PmEC01597 | 211 | 21-22 | 21.9 | NLS (144-176) | Nucleus, nucleolus, plasma membrane |
| PmEC01742 | 78 | 22-23 | 6.0 |  | Nucleus and cytosol |
| PmEC01936 | 221 | 20-21 | 21.6 |  | No accumulation |
| PmEC01984 | 121 | 23-24 | 10.7 |  | Nucleus and cytosol |
| PmEC02017 | 130 | 17-18 | 12.1 |  | Nucleus and cytosol |
| PmEC02274 | 86 | 18-19 | 7.1 |  | Nucleus and cytosol |
| PmEC02331 | 222 | 19-20 | 21.8 |  | Nucleus and cytosol |
| PmEC02451 | 150 | 17-18 | 14.2 |  | No accumulation |
| PmEC02707 | 95 | 17-18 | 8.7 |  | Nucleus and cytosol |
| PmEC02872 | 122 | 21-22 | 10.5 |  | Nucleus and cytosol |
| PmEC02890 | 180 | 23-24 | 17.3 |  | Nucleus and cytosol |
| PmEC02905 | 101 | 18-19 | 9.7 |  | Nucleus and cytosol |
| PmEC03053 | 299 | 23-24 | 30.7 |  | Nucleus and cytosol |
| PmEC03153 | 134 | 28-29 | 11.7 | Chloroplast (17-43) | Nucleus and cytosol |
| PmEC03234 | 142 | 20-21 | 13.2 |  | Nucleus and cytosol |
| PmEC03436 | 179 | 18-19 | 16.6 |  | Cytosol |
| PmEC03476 | 68 | 16-17 | 5.4 |  | Nucleus and cytosol |
| PmEC03493 | 212 | 24-25 | 21.7 | Mitochondria (52-73) | Nucleus, sub-nuclear, cytosol |
| PmEC03629 | 95 | 19-20 | 8.4 |  | Nucleus and cytosol |
| PmEC03706 | 219 | 19-20 | 21.2 | Chloroplast (29-56)<br>NLS (131-134) | Cytosol |
| PmEC03792 | 96 | 33-34 | 7.5 | NLS (8-30) | Nucleus and nucleolus |
| PmEC04014 | 191 | 20-21 | 18.9 |  | Cytosol and cytosolic aggregates |
| PmEC04128 | 281 | 15-16 | 30.4 |  | Nucleus and cytosol |
| PmEC04129 | 165 | 18-19 | 16.0 |  | Nucleus and cytosol |
| PmEC04322 | 205 | 16-17 | 20.5 |  | Nucleus and cytosol |
| PmEC04573 | 159 | 18-19 | 15.3 | Chloroplast (60-88) | Nucleus, cytosol, chloroplasts |
| PmEC05555 | 237 | 18-19 | 23.6 |  | Nucleus and cytosol |
| PmEC05617 | 143 | 21-22 | 13.9 | NLS (113-121) | Cytosol |
| PmEC06216 | 141 | 23-24 | 13.1 |  | No accumulation |
| PmEC06656 | 193 | 18-19 | 19.2 |  | Nucleus and cytosol |
| PmEC06699 | 138 | 19-20 | 13.0 |  | Nucleus and cytosol |
| PmEC06759 | 207 | 23-24 | 21.3 |  | Nucleus and cytosol |
| PmEC07010 | 197 | 21-22 | 19.1 |  | Nucleus and cytosol |

a. Data mined from Telenko et al., (2020).
b. Numbers indicate the length of amino acids including the predicted signal peptide.
c. Predicted signal peptide. Numbers indicate amino acid positions. Signal peptide predictions were performed using SignalP (v6.0).
d. Predicted molecular weight (in kilodaltons; kDa) of the mature protein without the signal peptide.
e. Subcellular localization as predicted by LOCALIZER (v1.0). Numbers within the parentheses indicate the amino acid positions within the predicted transit peptide sequence. NLS; nuclear localization signal.
f. Subcellular localization patterns in *Nicotiana benthamiana* epidermal cells determined in this study.

### Generation of plant expression constructs

All constructs in this study were generated using a modified multisite Gateway cloning system (Invitrogen). The AtUBQ10-NLS:mCherry, AtFLS2:mCherry, RbcS-TP:mCherry, AtFIB2:mCherry, 3xHA:sYFP (free sYFP), and 3xHA:mCherry (free mCherry) constructs have been described previously (Denne et al., 2021; Gu et al., 2011; Helm et al., 2019; Nelson et al., 2007; Qi et al., 2012; Robin et al., 2018).

A commercial gene synthesis service (Azenta Life Sciences, South Plainfield, New Jersey) was used to synthesize the open reading frames (ORFs) of each *P. maydis* effector candidate (PmEC) (without the signal peptide) with codon optimization for plant expression. The *attL*1 and *attL*4 Gateway sequences were added to the 5’ and 3’ ends, respectively, of each *P. maydis* candidate effector to generate Gateway-compatible DONR clones. The resulting sequences were inserted into the plasmid vector pUC57 by the service provider. We designated the resulting constructs pDONR(L1-L4):PmEC.

To generate the PmEC-sYFP protein fusions, the pDONR(L1-L4):PmEC constructs were mixed with pBSDONR(L4r-L2):sYFP (Qi et al., 2012), and the Gateway-compatible expression vector pEG100 (Earley et al., 2006), which places the transgene under control of the 35S promoter. All plasmids were recombined by the addition of LR Clonase II (Invitrogen) and were incubated overnight at 25°C following the manufacturer’s instructions. The resulting constructs were transformed into *Agrobacterium tumefaciens* GV3101 (pMP90) by electroporation and subsequently used for transient expression in *Nicotiana benthamiana*.

### Agrobacterium-mediated transient protein expression in *Nicotiana benthamiana*

The CaMV 35S-driven constructs described above were mobilized into *Agrobacterium tumefaciens* GV3101 (pMP90) and grown on LB medium plates containing 25 μg of gentamicin sulfate and 50 μg of kanamycin per milliliter for 2 days at 30°C. Cultures were prepared in liquid media (10 ml) supplemented with the appropriate antibiotics and were shaken overnight at 30°C at 225 rpm on an orbital shaker. Following overnight incubation, the cells were pelleted by either centrifuging at 3,600 rpm for 3 minutes or 3,000 rpm for 8 minutes at room temperature. The bacterial pellet was then resuspended in either 10 mM MgCl_2_ or induction medium (10 mM MES, pH 5.5, 3% sucrose), adjusted to an optical density at 600 nm (OD_600_) of 0.4 (final concentration for each strain in a mixture), and incubated with either 100 or 200 μM acetosyringone (Sigma-Aldrich) for 3-4 hours at room temperature with constant shaking. Bacterial suspensions were mixed in equal ratios (1:1) and infiltrated into the underside of 3- to 4-week-old *Nicotiana benthamiana* leaves with a needleless syringe. Leaf samples were collected 24 hours after agroinfiltration for immunoblot analyses or confocal microscopy.

### Protein extraction and immunoblot analyses

*N. benthamiana* leaf samples (0.5g) were collected at 24 hrs following agroinfiltration and flash frozen in liquid nitrogen. Tissue was homogenized with protein extraction buffer (50 mM Tris-HCl [pH 7.5], 150 mM NaCl, 0.1% Nonidet P-40, 1% 2,2’- dipyridyl disulfide [DPDS] and 1% protease inhibitor cocktail) (Sigma Aldrich). Homogenates were briefly mixed and centrifuged twice at 10,000 x g for 10 min at 4°C to pellet cell debris. Total protein lysates were combined with 4X Laemmli Sample Buffer (277.8 mM Tris-HCl [pH 6.8], 4.4% LDS, 44.4% (v/v) glycerol, 0.02% bromophenol blue, and 10% β-mercaptoethanol), and the mixtures were boiled at 95°C for 10 min. Protein samples were separated on 4-20% Tris-glycine stain-free polyacrylamide gels (Bio-Rad) at 175 V for 1 hr in 1X Tris/glycine/SDS running buffer. Total proteins were transferred to nitrocellulose membranes (GE Water and Process Technologies) at 100 V for one hour. Ponceau staining was used to confirm equal loading and transfer of total protein samples. Membranes were washed with 1X Tris-buffered saline (50 mM Tris-HCl, 150 mM NaCl [pH 6.8]) solution containing 0.1% Tween20 (TBST) and incubated with 5% Difco skim milk for 1 hr at room temperature or overnight at 4°C with gentle shaking. Proteins were subsequently detected with horseradish peroxidase (HRP)-conjugated anti-GFP antibody (1:5,000) (Miltenyi Biotec) for 1 hr at room temperature with gentle shaking. Following antibody incubation, membranes were washed at least three times for 15 minutes in 1x TBST solution. Protein bands were imaged using equal parts of Clarity Western ECL substrate peroxide solution and luminol/enhancer (BioRad) solution (Thermo Scientific), with incubation at room temperature for 5 minutes. Immunoblots were developed using X-ray film.

### Confocal microscopy

Live-cell imaging of *N. benthamiana* epidermal cells was performed 24 hours following agroinfiltration using a Zeiss LSM880 Axio Examiner upright confocal microscope as described previously (Denne et al., 2021). Briefly, *N. benthamiana* leaf sections were excised and mounted in sterile water between a slide and a coverslip (adaxial surface toward the objective) and subsequently imaged using a Plan Apochromat 20x/0.8 objective, pinhole 1.0 AU. For plasmolysis, leaf sections were prepared as described above, submerged in 0.8 M mannitol solution for 20 minutes, and imaged shortly thereafter. The sYFP protein fusions were excited using a 514-nm argon laser and fluorescence was detected between 517-562 nm. Fluorescence from the mCherry-tagged constructs was excited with a 561-nm helium-neon laser and detected between 565-669 nm. All confocal micrographs were captured on the Zeiss LSM880 upright confocal microscope and processed using the Zeiss Zen Blue Lite program (Carl Zeiss Microscopy, USA).

## RESULTS

### *In silico* selection and generation of super Yellow Fluorescent Protein (sYFP)- tagged *P. maydis* effector candidate proteins (PmECs) from Indiana isolate PM-01

To identify promising secreted effector candidates, we leveraged effector predictions previously generated by Telenko et al. (2020). These authors identified 59 *P. maydis* proteins that contain effector-like characteristics as determined by EffectorP (v2.0). To further refine the previous analyses by Telenko et al. (2020), we mined through the 59 *P. maydis* effector candidates (PmECs) and selected proteins that fulfilled more selective criteria: i) size fewer than 300 amino acids; ii) presence of a signal peptide (as predicted by SignalP v6.0); and iii) lack of a transmembrane domain (as predicted by TMHMM v2.0). We next leveraged EffectorP (v3.0) to identify putative effector proteins from this narrowed set of proteins. Among the 59 potential effectors originally identified by Telenko et al. (2020), 40 contain effector-like protein characteristics that fulfilled these more selective criteria (Figure 1A). The selected effector candidates ranged in size from 55 to 299 amino acids (Table 1). We next employed LOCALIZER (v1.0) to identify predicted nuclear localization signals (NLS), chloroplast transit peptides, or mitochondrial targeting sequences. As shown in Table 1, many of the candidates are not predicted to target specific subcellular compartments. However, several PmECs encode predicted nuclear localization signals including PmEC01597, PmEC03792, and PmEC05617 (Table 1). Two effector candidates, PmEC03153 and PmEC04573, contain predicted chloroplast transit peptides, and a mitochondrial targeting sequence was identified in PmEC03493 (Table 1). Intriguingly, PmEC00848 encoded both a chloroplast transit peptide and a mitochondrial-targeting sequence and PmEC03706 encoded both a chloroplast transit peptide and NLS, suggesting these proteins may localize to multiple subcellular compartments (Table 1).

To investigate the subcellular compartments targeted by the *P. maydis* effector candidates, the predicted open reading frames (ORFs) of each of the 40 putative effectors, without the predicted signal peptides, were synthesized and fused to the N terminus of super Yellow Fluorescent Protein (PmEC:sYFP) (Figure 1B). The resulting constructs were recombined into the plant expression binary vector pEarleyGate100 (pEG100) (Earley et al., 2006), which places the candidate effectors downstream of a 35S promoter. The resulting constructs were inserted into *Agrobacterium tumefaciens* for subsequent *Nicotiana benthamiana*-based heterologous expression assays (Figure 1C-E).

### Candidate *P. maydis* effector-fluorescent protein fusions accumulate protein *in planta*

Prior to assessing the subcellular localization of the *P. maydis* effector-fluorescent protein fusions, we tested whether these effectors accumulate protein in dicot leaf cells. This was accomplished by transiently expressing each of the fusion proteins in *N. benthamiana* using *Agrobacterium*-mediated infiltration (agroinfiltration). Immunoblot analyses revealed that, of the 40 candidate effectors screened, 37 accumulated at detectable levels when transiently expressed in *N. benthamiana* (Figure 2). Three putative effectors (PmEC01936, PmEC02451 and PmEC06216) consistently failed to express detectable protein suggesting these fusion proteins do not accumulate when transiently expressed in *N. benthamiana* leaf cells (Figure 2). Though most of the fusion proteins accumulated at the expected molecular weight, several putative effectors accumulated protein at lower molecular weights than predicted, suggesting post-translational modifications (Figure 2). As most *P. maydis* effector-fluorescent protein fusions accumulated protein, we conclude that *N. benthamiana* is an appropriate surrogate plant system and can thus be used to investigate the subcellular localization patterns of *P. maydis* effector-fluorescent protein fusions. Based on these data, we discarded the effector candidates with insufficient protein expression (PmEC01936, PmEC02451 and PmEC06216) and retained the remaining candidate effectors for further *in planta* analyses.

**Figure 2.**
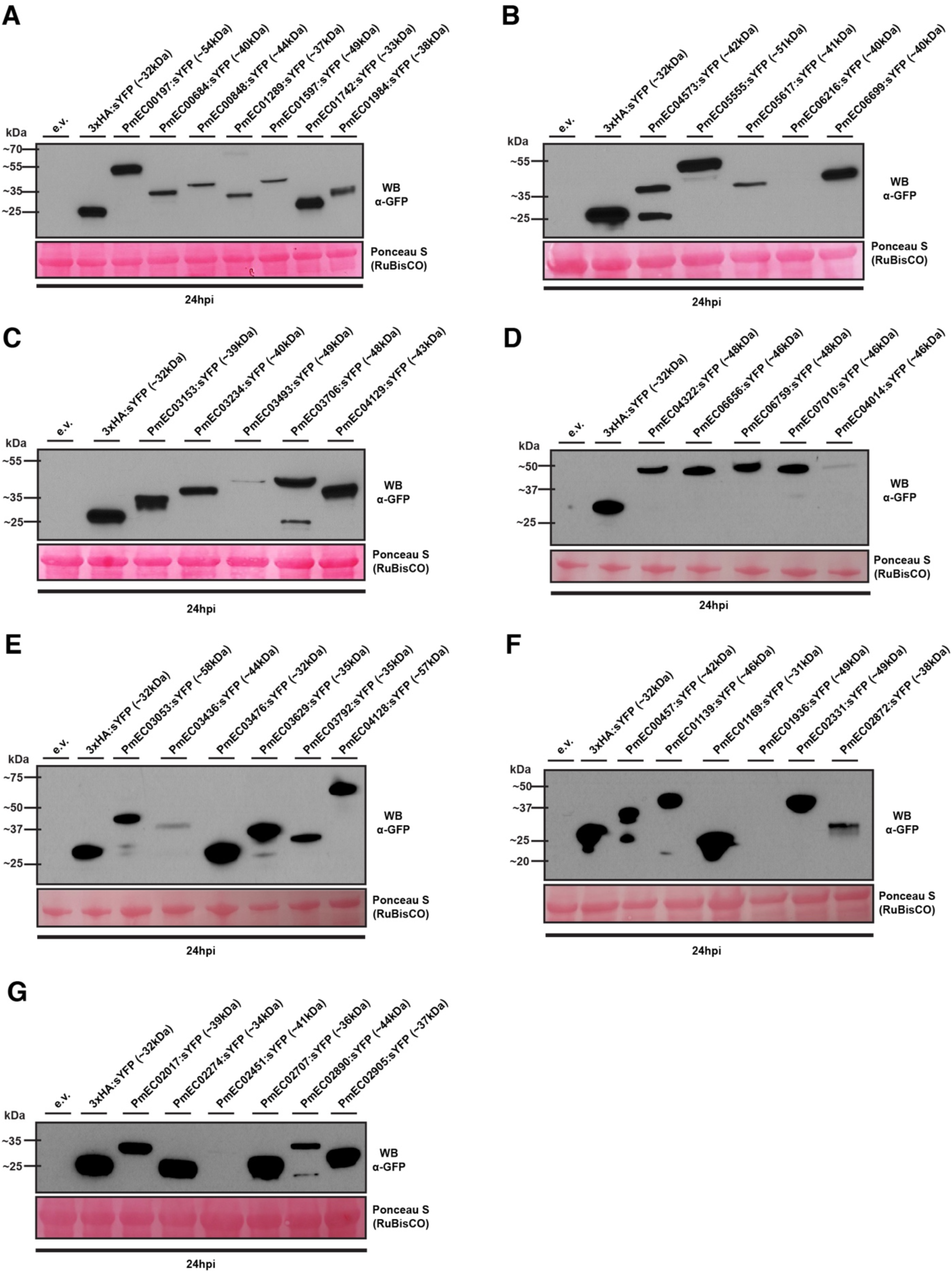
Immunoblot analyses of the *Phyllachora maydis* candidate effector-fluorescent fusion proteins. For panels A-G), the indicated constructs were transiently expressed in 3-week-old *Nicotiana benthamiana* using agroinfiltration. Total protein was extracted 24 hours post-agroinfiltration and immunoblotted with horseradish peroxidase (HRP)-conjugated anti-GFP antibodies. Free sYFP (3xHA:sYFP) and empty vector (e.v.) were included as controls. Ponceau staining of the RuBisCO large subunit was used as a loading control. For each protein, the theoretical protein size is indicated in parentheses (in kilodaltons; kDa). Three independent experiments were performed with similar results. The results of only one experiment are shown.

### The majority of *P. maydis* effector candidate-fluorescent protein fusions localize to the nucleus and cytosol

Live-cell imaging of epidermal cells using laser-scanning confocal microscopy revealed that among the 37 sYFP-tagged PmECs that accumulate protein, 29 showed subcellular distribution patterns in the nucleo-cytosol and were indistinguishable from the free sYFP control (Supplemental Figure 1; Table 1). Furthermore, fluorescence signal from five sYFP-tagged derivatives, PmEC03436, PmEC03493, PmEC03706, PmEC04014, and PmEC05617, predominantly accumulated in the cytosol (Supplemental Figure 2; Table 1). Interestingly, PmEC03436:sYFP signal labeled punctate bodies on the cell periphery, and PmEC04014:sYFP signal was observed in irregular, cytosolic aggregates (Supplemental Figure 2). In addition to localizing in the cytosol, PmEC03493:sYFP accumulated in the nucleus as well as sub-nuclear structures (Supplemental Figure 2). The remaining effector-fluorescent protein fusions preferentially localized to specific subcellular compartments within the plant cells.

### The *P. maydis* effector candidate PmEC01597 localizes to multiple subcellular compartments

PmEC01597 encodes a predicted nuclear localization signal (NLS) at its C terminus as predicted by LOCALIZER (Sperschneider et al., 2017) (Table 1; Figure 3A) and a nucleolar targeting signal (NoLS) as predicted by NoD (http://www.compbio.dundee.ac.uk/www-nod/; Scott et al., 2011), suggesting this effector candidate localizes to both the nucleus and nucleolus. To confirm the specific localization of PmEC01597 to these subcellular compartments, we co-expressed PmEC01597:sYFP with mCherry-tagged AtUBQ10-NLS or AtFIB2, Arabidopsis proteins known to localize to the nucleus and nucleolus, respectively (Nelson et al., 2007; Robin et al., 2018). As predicted, the PmEC01597:sYFP fluorescence signal consistently co-localized with both the AtUBQ10-NLS:mCherry and AtFIB2:mCherry fluorescence signals, demonstrating that PmEC01597:sYFP accumulates in the nucleus and nucleolus (Figure 3C-D). Furthermore, PmEC01597:sYFP also overlapped fluorescence signals (Figure 3E) with mCherry-tagged FLS2 (FLS2:mCherry), an Arabidopsis protein known to localize on the plasma membrane (Helm et al., 2019). Plasmolysis of *N. benthamiana* epidermal cells expressing PmEC01597:sYFP and AtFLS2:mCherry revealed separation of the plasma membrane from the cell wall, further supporting plasma-membrane localization of PmEC01597:sYFP (Supplemental Figure 3). Collectively, these data demonstrate that PmEC01597 targets multiple subcellular compartments when expressed in *N. benthamiana*.

**Figure 3.**
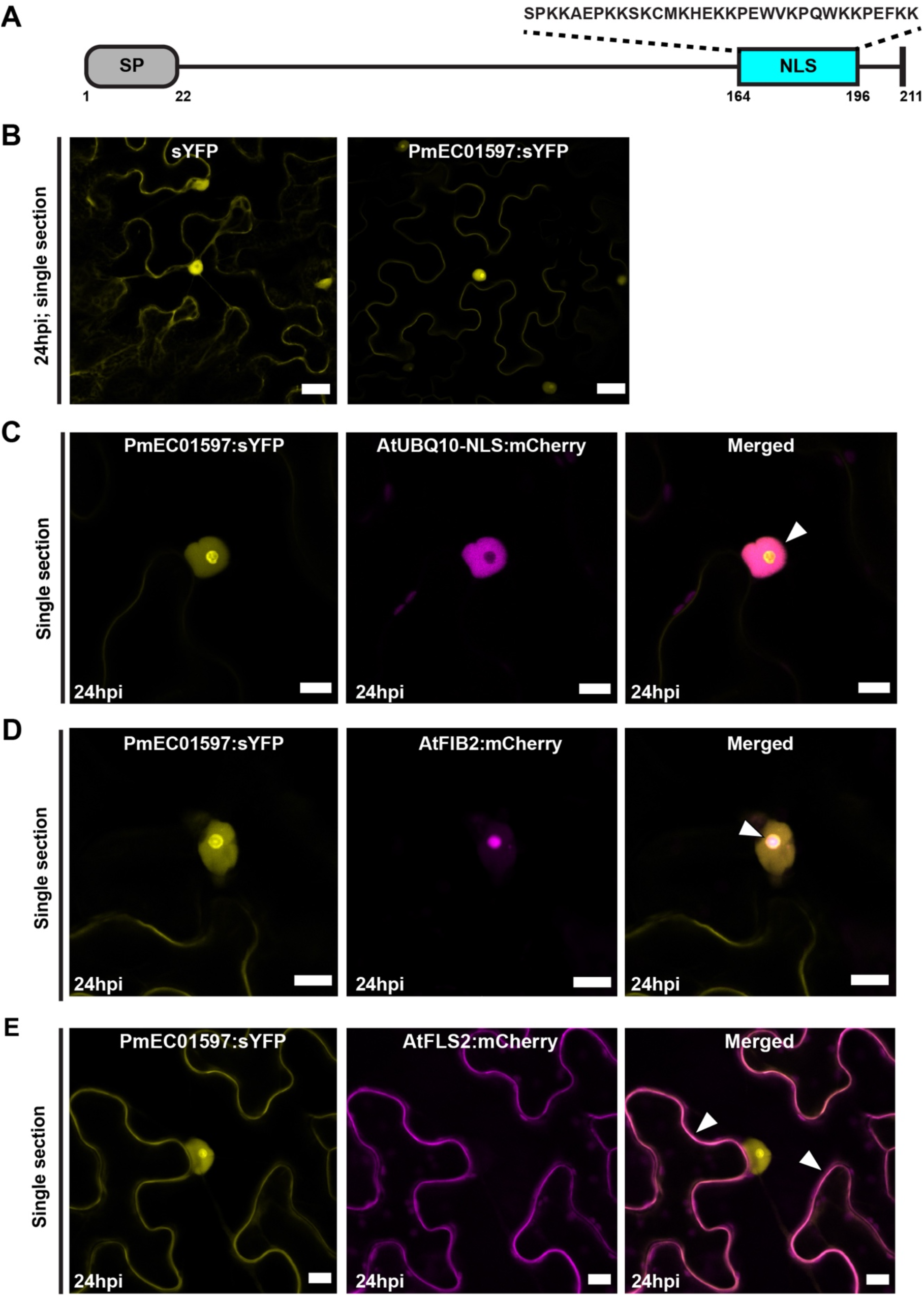
The PmEC01597-fluorescent protein fusions accumulate within the nucleus, nucleolus, and on the plasma membrane in *N. benthamiana*. **A)** Schematic representation of the *P. maydis* PmEC01597 candidate effector including the nuclear localization signal (NLS). Numbers beneath the schematic illustration represent amino acid positions. **B)** Live-cell imaging of PmEC01597:sYFP in *N. benthamiana* epidermal cells. Free sYFP (left panel) was included as a reference for nucleo-cytoplasmic distribution. The scale bar shown represents 20 µM. **C)** Live-cell imaging of PmEC01597:sYFP and AtUBQ10-NLS:mCherry in *N. benthamiana* leaf pavement cells. The scale bars shown represent 10 µM. **D)** PmEC01597:sYFP fusion proteins localize to the nucleolus in *N. benthamiana*. mCherry-tagged Arabidopsis FIB2 was included as a reference for nucleolus localization. The scale bars shown represent 10 µM. **E)** PmEC01597:sYFP fusion proteins localize on the plasma membrane in *N. benthamiana*. mCherry-tagged Arabidopsis FLS2 was included as a reference for plasma membrane localization. For panels B-E), all confocal micrographs shown are of single optical sections and white arrowheads indicate overlapping sYFP and mCherry fluorescence signals.

### The *P. maydis* effector candidate PmEC03792 is imported into the nucleus and nucleolus

Fungal pathogens have been shown to express and translocate effectors that preferentially accumulate protein within the host nucleus (Kemen et al., 2005; Petre et al., 2015). We, therefore, investigated whether any of the *P. maydis* effector candidates specifically targeted the nucleus. Analysis of the PmEC03792 protein sequence revealed a NLS motif at its N terminus (aa 8-30) as well as a nucleolar-targeting sequence (aa 4-35), suggesting that it may localize to the nucleus as well as the nucleolus (Sperschneider et al., 2017; Scott et al., 2011) (Table 1). To test our hypothesis, we transiently expressed the PmEC03792:sYFP protein fusion and assessed the subcellular localization pattern in *N. benthamiana* epidermal cells. Live-cell imaging using laser scanning confocal microscopy showed that PmEC03792 preferentially localized to subcellular compartments resembling the nucleus and nucleolus, whereas free sYFP predominantly localized to the cytoplasm and the nucleus (Figure 4). To confirm PmEC03792 is indeed localized to the nucleus and nucleolus, we transiently co-expressed PmEC03792:sYFP with either mCherry-tagged AtUBQ10-NLS or AtFIB2. Consistent with our hypothesis, live-cell imaging revealed that the PmEC03792:sYFP fluorescence signal co-localized with both the AtUBQ10-NLS:mCherry and AtFIB2:mCherry fluorescence signals, demonstrating that PmEC03792:sYFP accumulates in the nucleus and nucleolus (Figure 4).

**Figure 4.**
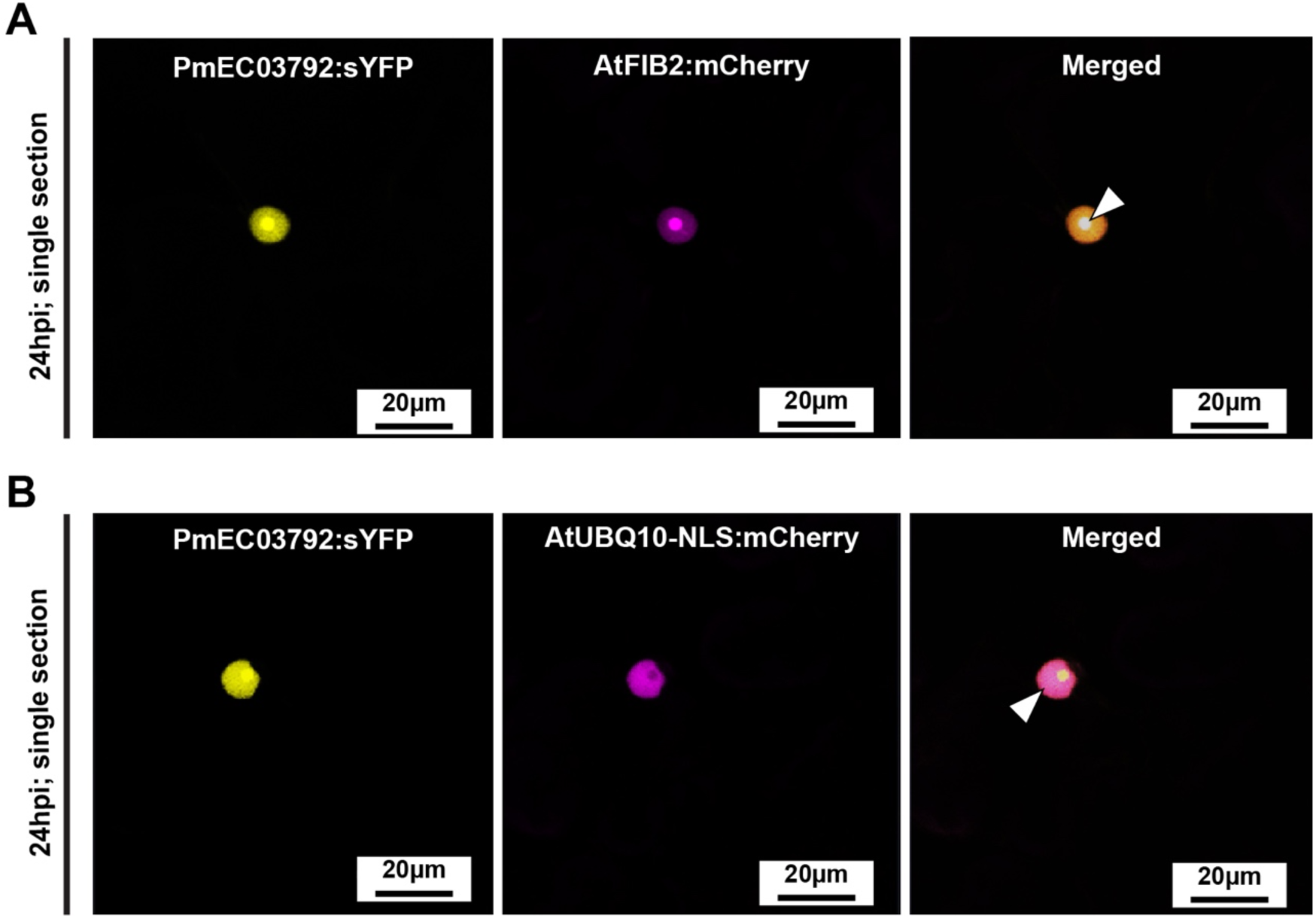
The PmEC03792-fluorescent protein fusion preferentially localizes to the nucleolus and nucleus in *N. benthamiana*. Live-cell imaging of *N. benthamiana* leaf pavement cells expressing **A)** PmEC03792:sYFP and AtFIB2:mCherry, and **(B)** PmEC03792:sYFP and AtUBQ10-NLS:mCherry. White arrowheads indicate overlapping of sYFP and mCherry fluorescence signals. Images are single optical sections. For panels A and B), protein fusions were expressed in *N. benthamiana* using agroinfiltration and live-cell imaging was performed 24 hours following agroinfiltration.

### PmEC04573 targets the chloroplasts

Chloroplasts often have an essential role in coordinating an effective plant immune response against pathogens and, as such, are often targeted by proteinaceous effectors from filamentous fungal pathogens (Littlejohn et al., 2021). Indeed, several of the *P. maydis* effector candidates encode predicted chloroplast transit peptide (cTP) sequences (Table 1), suggesting these effector candidates may target host chloroplasts. We, therefore, investigated whether any of the *P. maydis* effector-fluorescent protein fusions localized to these subcellular compartments. Intriguingly, fluorescence from the PmEC04573:sYFP-fluorescent protein fusion was consistently detected in organelles resembling chloroplasts as well as the nucleo-cytosol (Figure 5A-D). To test whether PmEC04573:sYFP is indeed chloroplast-localized, we co-expressed PmEC04573:sYFP with RbcS-TP:mCherry, a subcellular marker for plastids (Nelson et al., 2007). Consistent with our hypothesis, PmEC04573:sYFP fluorescence signal overlapped with RbcS-TP:mCherry on chloroplasts as well as stromules (stroma-containing tubules), confirming PmEC04573:sYFP does indeed accumulate on chloroplasts (Figure 5B-C).

**Figure 5.**
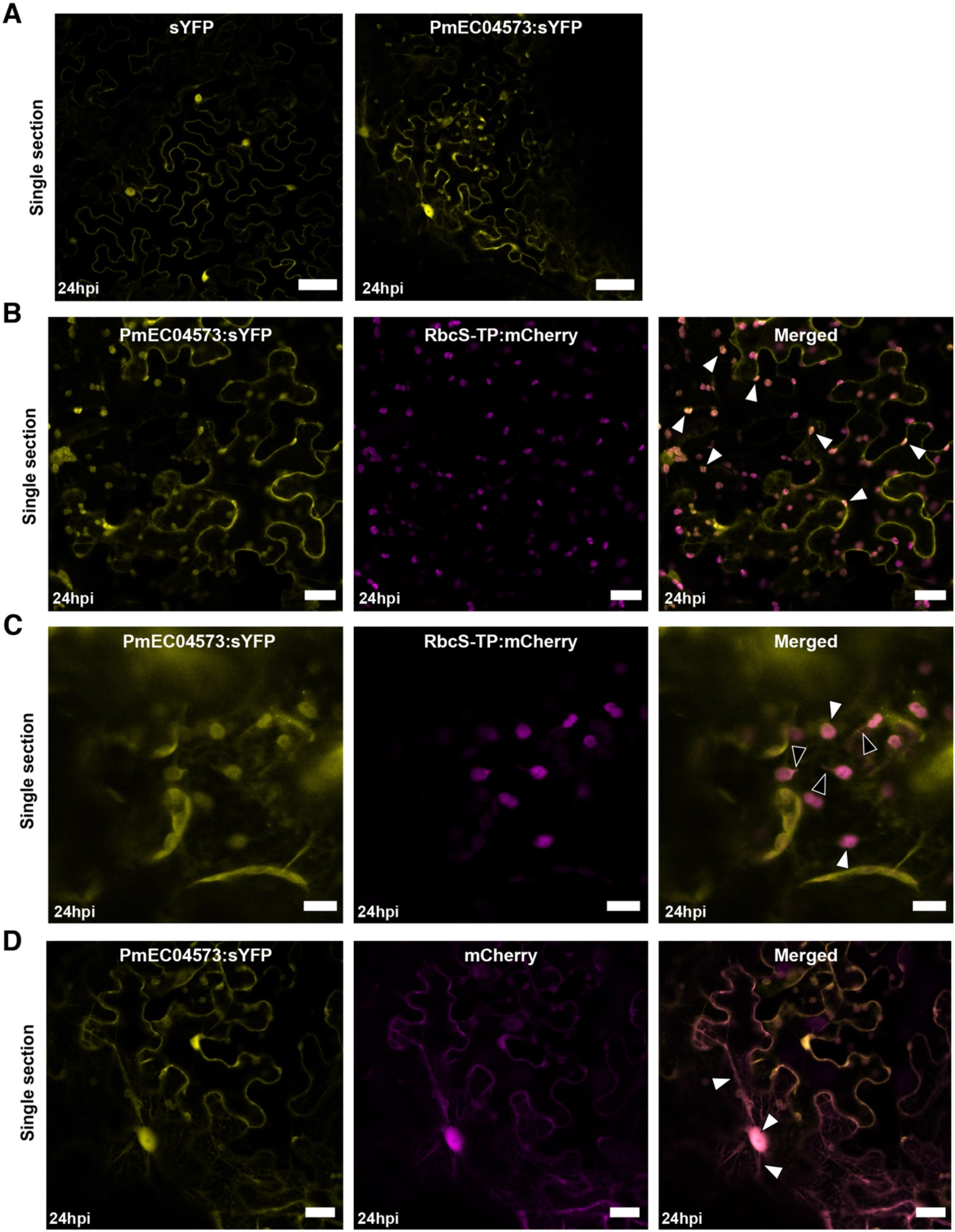
PmEC04573-fluorescent protein fusions accumulate on chloroplasts. **A)** Live-cell imaging of *P. maydis* PmEC04573:sYFP in *N. benthamiana* epidermal cells. Confocal micrographs are single optical sections. Free sYFP (left panel) was included as a reference for nucleo-cytoplasmic distribution. The scale bar shown represents 50 µM. **B-C)** Live-cell imaging of PmEC04573:sYFP and RbcS-TP:mCherry (plastid marker) in *N. benthamiana* leaf pavement cells. The scale bars shown represent 20 µM in panel b and 10 µM in panel c. **D)** Live-cell imaging of PmEC04573:sYFP and free mCherry in *N. benthamiana* leaf pavement cells. The scale bars shown represent 20 µM. For panels A-D), protein fusions were expressed in *N. benthamiana* using agroinfiltration and live-cell imaging was performed 24 hours following agroinfiltration. All confocal micrographs shown are of single optical sections and white arrowheads indicate overlapping sYFP and mCherry fluorescence signals. Black arrowheads indicate stromules.

Interestingly, immunoblot analyses with the PmEC04573:sYFP protein fusion consistently revealed two distinct protein products; the larger protein product was near the predicted molecular weight while the smaller protein band coincided with free sYFP, suggesting cleavage of sYFP from PmEC04573 had occurred (Figure 2; Supplemental Figure 4). To exclude that the chloroplast localization observed with PmEC04573:sYFP was an artifact caused by free sYFP, we coexpressed sYFP with RbcS-TP:mCherry in *N. benthamiana* leaf cells. As predicted, sYFP fluorescence signal was not observed on chloroplasts, even when the sYFP signal was saturated (Supplemental Figure 5). These protein expression data, coupled with the observation that free sYFP did not localize to chloroplasts, suggests that the nucleo-cytosol localization observed with PmEC04573:sYFP (Figure 5D) may be attributed to processed sYFP diffusing throughout the nucleoplasm and cytoplasm.

## DISCUSSION

Recent sequencing of the *Phyllachora maydis* genome revealed that this fungal pathogen encodes putatively secreted effector candidate proteins (Telenko et al., 2020). However, our general understanding of host cell compartments targeted by the *P. maydis* effector repertoire is limited (Helm et al., 2022). In this study, we leveraged the availability of the *P. maydis* genome and refined previous effector predictions to investigate the subcellular compartments targeted by candidate effector proteins using a *Nicotiana benthamiana*-based heterologous expression system (Figure 1). We found that 37 of the 40 putative effectors accumulated detectable protein *in planta* (Figure 2). Among the 37 *P. maydis* effector-fluorescent protein fusions tested, 29 displayed a nucleo-cytoplasmic distribution that was indistinguishable from the free sYFP control (Supplemental Figure 1) and five predominantly localized to the cytosol (Supplemental Figure 2). One effector candidate, PmEC01597, localized to multiple subcellular compartments including the nucleus, nucleolus, and plasma membrane (Figure 3), while PmEC03792 was specifically imported into both the nucleus and nucleolus with no observable cytoplasmic stranding (Figure 4). Another putative effector, PmEC04573, consistently localized to chloroplasts as well as stromules (Figure 5).

Collectively, our data suggest that candidate effector proteins from *P. maydis* localize to distinct subcellular compartments and may associate with host proteins in these locations. It should be acknowledged that our approach relied on fusing a large fluorophore to the C terminus of each candidate effector as well as overexpression in a heterologous model plant. Furthermore, the selected putative effectors are predicted to encode signal peptides and are thus likely to be secreted; however, there is no direct evidence these proteins are translocated into host cells. Nevertheless, the observation that some PmECs accumulated protein and were targeted to specific subcellular locations within leaf cells suggests that these proteins are *bona fide* cytoplasmic effectors. Hence, knowledge of the subcellular localization patterns of the *P. maydis* effector repertoire will be informative in the identification of the host proteins they target as well as the cellular pathways they alter. Determining whether any of the *P. maydis* effector candidates have a functional role in manipulating host immune responses as well as their potential host targets in maize is a focus for future investigations.

One effector candidate, PmEC01597, consistently localized to multiple subcellular compartments including the nucleus, nucleolus, and plasma membrane (Figure 3). Though it is unclear of the functional significance of PmEC01597 localization to the nucleolus, effectors from fungi and oomycetes have been shown to target this subcellular compartment (Lorrain et al., 2018). A candidate effector from the poplar leaf rust pathogen (*Melamspora larici-populina*), termed Mlp124478, encodes a predicted nuclear localization peptide sequence, and accumulated in the nucleus and nucleolus of *N. benthamiana* epidermal cells (Ahmed et al., 2018; Petre et al., 2015). Furthermore, transgenic Arabidopsis constitutively expressing Mlp124478 displayed altered leaf morphology as well as repressed defense gene expression (Ahmed et al., 2018). Intriguingly, Chip-PCR analyses revealed that this candidate effector associates with the TGA1a-binding DNA sequence, suggesting Mlp124478 binds the TGA1a promoter region and represses expression of defense genes (Ahmed et al., 2018). Though the biological significance of PmEC01597 accumulating in the nucleolus remains to be investigated, we hypothesize this putative effector may interfere with host cell transcriptional machinery of ribosomal RNA (rRNA) genes or processing of ribosomal RNA synthesis.

In addition to targeting the nucleus and nucleolus, PmEC01597 consistently localized on the plasma membrane when transiently expressed in *N. benthamiana* epidermal cells (Figure 3). Proteinaceous effectors from filamentous phytopathogens have indeed been shown to target the plasma membrane where they often modulate host immune responses (Fabro, 2022; Lorrain et al., 2018). For example, work by Gaouar and colleagues (2016) showed that a different putative effector from poplar leaf rust, Mlp124202, localized on the plasma membrane when transiently expressed in *N. benthamiana* and in stable transgenic Arabidopsis. Consistent with the subcellular localization, yeast two-hybrid assays revealed that Mlp124202 associated with plasma membrane-localized synaptotagmin A (SYTA; At2g20990), suggesting this poplar leaf rust effector may have a functional role in modulating vesicle-mediated trafficking. Furthermore, the *Phytophthora sojae*-secreted effector, Avh240, preferentially accumulated on the host plasma membrane where it associated with and inhibited secretion of the soybean aspartic protease, GmAP1, thereby suppressing host immune responses (Guo et al., 2019). Murphy and colleagues (2018) showed that the *Phytophthora infestans* effector, Pi17316, interacts directly with VASCULAR HIGHWAY1-interacting kinase from potato (StVIK). Importantly, transgenic overexpression of StVIK in potato enhanced *Phytophthora infestans* virulence and colonization, demonstrating StVIK functions at least in part as a susceptibility factor (Murphy et al., 2018). We, therefore, predict PmEC01597 associates with host proteins on the plasma membrane to modulate host immune responses, similar to those of other plasma membrane-localized effectors from fungal and oomycete pathogens. Hence, future functional characterization of PmEC01597 should prioritize identifying host proteins from maize that interact with this putative effector.

Numerous filamentous phytopathogens often express and translocate effectors inside host cells where they specifically localize to host nuclear compartments and manipulate host immune responses (Caillaud et al., 2012; Schornack et al., 2010; Stam et al., 2013). Here, we show one putative effector, PmEC03792, was specifically targeted to the nucleus and nucleolus (Figure 4). Consistent with the subcellular localization pattern, PmEC03792 encoded a predicted NLS motif as well as a nucleolar-targeting sequence, suggesting this putative effector may manipulate host nucleolar functions. Indeed, numerous nuclear-localized fungal and oomycete effectors have been shown to disrupt many cellular processes by reprogramming transcriptional mechanisms to suppress host immune response. For example, AVR2, an effector from *Phytophthora infestans*, targets the nucleus and suppresses host immune responses through its association with a brassinosteroid-responsive bHLH transcription factor (Boch and Bonas, 2010; Turnbull et al., 2017). Furthermore, the *Colletotrichum graminicola* effector, CgEP1, specifically targets the host nucleus where it binds to chromatin, indicating that CgEP1 may modulate host transcription (Vargas et al., 2016). Moreover, the *Ustilago maydis* effector, See1, is localized to the host nucleus and interacts with the host SGT1 protein to reactivate DNA synthesis and cell division in infected leaves (Redkar et al., 2015). We, therefore, speculate nuclear-localized PmEC03792 may have a functional role in manipulating host transcription by targeting host proteins associated with nuclear compartments. Future functional characterization of PmEC03792 should prioritize identifying host proteins targeted by this candidate effector.

Our finding that the *P. maydis* putative effector PmEC04573 labels chloroplasts suggests that this fungal pathogen may target this organelle to modulate chloroplast-mediated host immune responses (Figure 5). Indeed, filamentous fungal pathogens have evolved intracellular effectors that localize to chloroplasts wherein they subvert chloroplast-derived immune responses (Littlejohn et al., 2021). For example, *Melampsora larici-populina* secretes several putative effectors, termed Chloroplast Targeting Proteins (CTP1, CTP2, and CTP3), that accumulate in the stroma of chloroplasts when transiently expressed in *N. benthamiana* (Petre et al., 2015; Petre et al., 2016). Importantly, CTP1, CTP2, and CTP3 encode predicted chloroplast transit peptides that are cleaved upon their translocation to chloroplasts, and which are necessary and sufficient for chloroplast localization (Petre et al., 2016). The observation that PmEC04573 also encodes a predicted chloroplast transit peptide sequence and localizes to chloroplasts suggests the cTP sequence may be necessary for chloroplast localization.

The wheat stripe rust pathogen *Puccinia striiformis* f. sp. *tritici* (Pst) has also been shown to express and translocate several effectors into host cells where they subsequently traffic to chloroplasts (Figueroa et al., 2021; Littlejohn et al., 2021). For example, Pst_12806 is a haustorium-specific effector that, when secreted, localizes to host chloroplasts, and interacts with the photosynthesis-related protein TaISP (Xu et al., 2019). Importantly, the direct association between Pst_12806 and TaISP suppresses chloroplast-derived immune responses and photosynthesis, thereby promoting pathogen growth (Xu et al., 2019). Furthermore, two additional wheat stripe rust effectors, Pst_4 and Pst_5, were recently shown to interact with TaISP and attenuate chloroplast-derived immune responses (Wang et al., 2021). However, unlike Pst_12806, Pst_4 and Pst_5 associate with TaISP in the cytoplasm and such interaction likely prevents TaISP trafficking to the chloroplast, thereby suppressing chloroplast-derived production of reactive oxygen species (Wang et al., 2021). We, therefore, predict the subcellular targeting of chloroplasts by *P. maydis* may be important for facilitating infection. Future work should focus on identifying host proteins from maize that associate with PmEC04573, and what effect such interactions have on facilitating *P. maydis* infection.

We leveraged the availability of the *P. maydis* genome generated using short-read sequencing technology to select the candidate effectors investigated in our study (Telenko et al., 2020). However, given the relatively low BUSCO (benchmarking sets of universal single-copy orthologs) score and the high percentage of repetitive sequences within the fungal genome, we speculate the current *P. maydis* genome assembly is incomplete. Hence, future work should aim to generate an improved *P. maydis* genome using both short- and long-read sequencing technologies as such an improved genome will likely identify additional *P. maydis* effector candidates.

In summary, we show the majority of putative effectors from *P. maydis* accumulate protein *in planta*, and several localize to specific plant cell compartments including the nucleus, nucleolus, plasma membrane, and chloroplasts. Our data provide valuable insights into the putative functions of the *P. maydis* effector candidates as well as the host processes potentially manipulated by this fungal pathogen. Lastly, our findings can be used to generate testable hypotheses for addressing the functional roles of *P. maydis* effectors during pathogenicity as well as identifying their host targets in maize.

## ACKNOWLEDGEMENTS

The authors thank Darcy Telenko (Purdue University) for providing sequences of the candidate effectors from *P. maydis* Indiana isolate PM-01, and Roger Innes (Indiana University) for providing the AtFLS2:mCherry, AtFIB2:mCherry, 3xHA:sYFP (free sYFP), and 3xHA:mCherry (free mCherry) constructs. The authors would like to thank the Purdue University Imaging Facility for access to the Zeiss LSM880 Axio Examiner upright confocal microscope. We also thank Morgan Carter and Martin Darino for insightful discussions and critical reading of the manuscript. The funding bodies had no role in designing the experiments, collecting the data, or writing the manuscript. All opinions expressed in this paper are the authors’ and do not necessarily reflect the policies and views of USDA, DOE, or ORAU/ORISE. USDA is an equal opportunity provider and employer.

## DATA AVAILABILITY STATEMENT

The data that support the findings of this study are available from the corresponding author upon request.

## CONFLICT OF INTEREST

The authors declare that they have no competing interests and that the research was conducted in the absence of any commercial or financial relationships that could be construed as a potential conflict of interest

## AUTHOR CONTRIBUTIONS

M.H. and R.S. conceived and designed the study. M.H., R.S., R.H., N.J., and A.M., performed the experiments. M.H., R.S., R.H., N.J., A.S.I-P, and S.B.G. analyzed the data. M.H. and R.S. drafted and wrote the manuscript. All authors edited the manuscript and approved the final version.

## SUPPLEMENTARY FIGURE LEGENDS

**Supplemental Figure 1.**
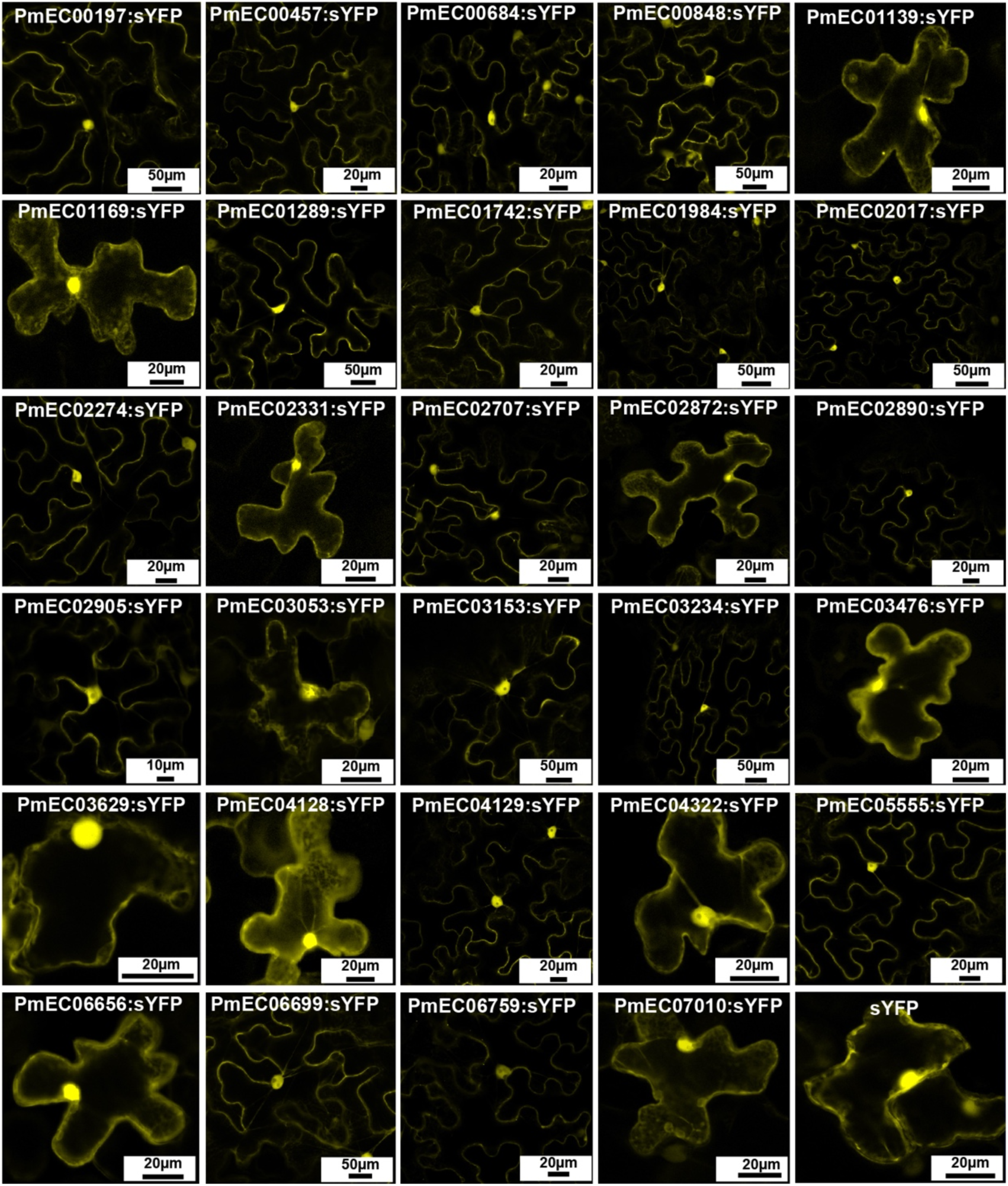
sYFP-tagged *P. maydis* effector candidate proteins localize to the nucleus and cytosol in *N. benthamiana*. The indicated constructs were transiently expressed in *N. benthamiana* and imaged using laser-scanning confocal microscopy 24 hours following agroinfiltration. Confocal micrographs shown are single optical sections. Free sYFP was included as a reference for nucleo-cytoplasmic distribution.

**Supplemental Figure 2.**
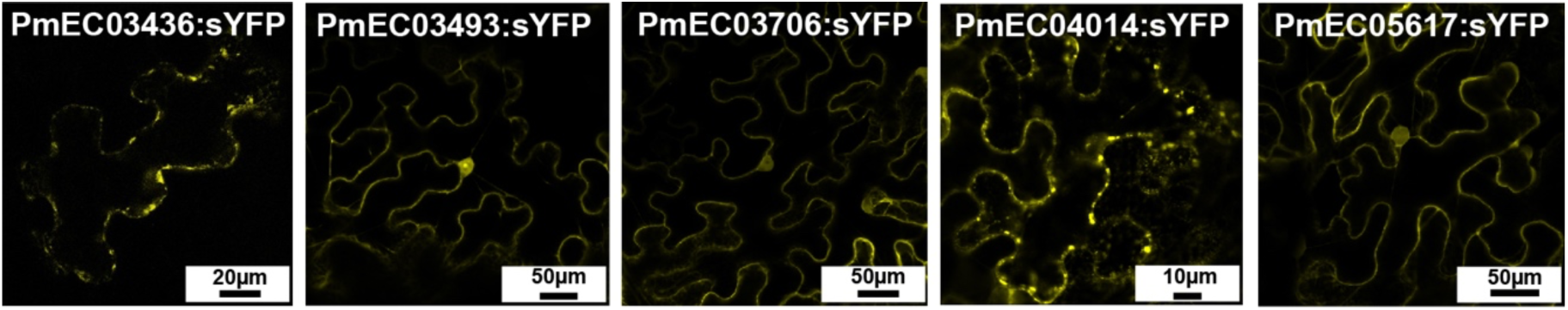
PmEC03436, PmEC03493, PmEC03706, PmEC04014, and PmEC05617 fluorescent protein fusions predominantly accumulate in the cytoplasm in *N. benthamiana*. Live-cell imaging of PmEC05617:sYFP and PmEC03706:sYFP in *N. benthamiana* epidermal cells. The indicated constructs were transiently expressed in *N. benthamiana* and imaged using laser-scanning confocal microscopy 24 hours following agroinfiltration. Confocal micrographs shown are single optical sections.

**Supplemental Figure 3.**
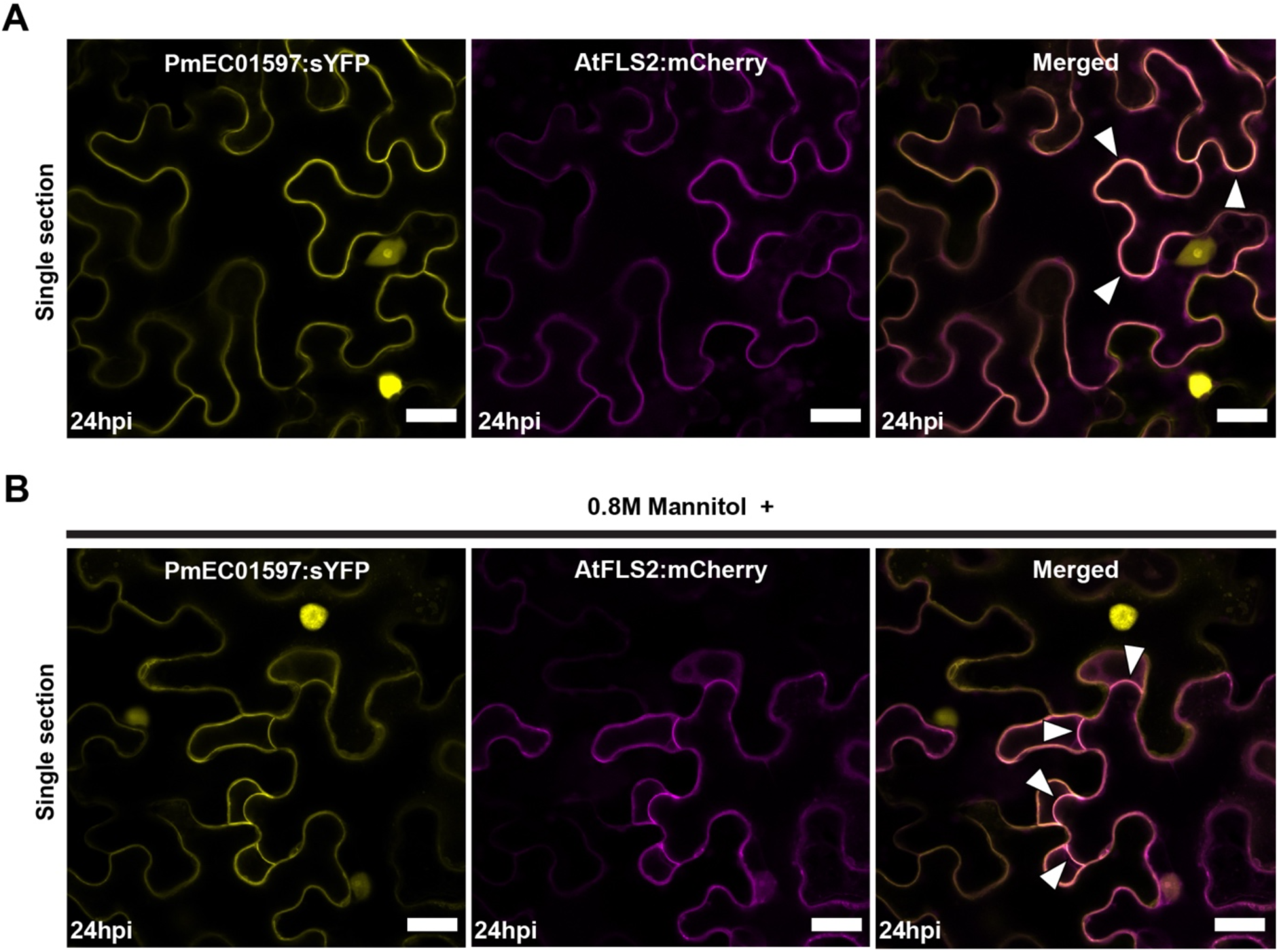
Plasmolysis of *N. benthamiana* epidermal cells expressing PmEC01597:sYFP and AtFLS2:mCherry fluorescent protein fusions reveal plasma membrane localization. **A)** PmEC01597:sYFP fusion proteins localize on the plasma membrane as indicated by co-localization with AtFLS2:mCherry. Images are of single optical sections. White arrowheads indicate overlapping sYFP and mCherry fluorescence signals. The scale bars shown represent 20 µM. **B)** Twenty-four hours following agroinfiltration, leaf sections of *N. benthamiana* expressing PmEC01597:sYFP and AtFLS2:mCherry were submerged in 0.8 M mannitol for 20 minutes to induce plasmolysis and live-cell imaging was performed shortly thereafter. White arrowheads indicate plasma membrane separation from the adjacent cell. The scale bar represents 20 µM.

**Supplemental Figure 4.**
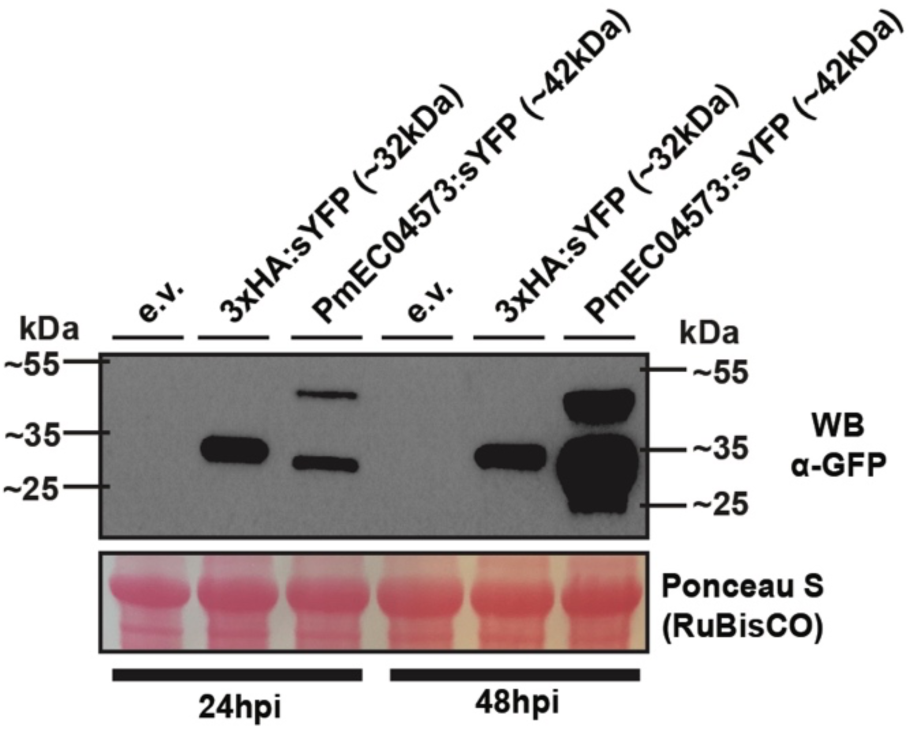
Immunoblot analyses of the PmEC04573:sYFP fusion proteins. The indicated constructs were transiently expressed in 3-week-old *N. benthamiana* using agroinfiltration. Total protein was extracted at 24 and 48 hours post-agroinfiltration and immunoblotted with HRP-conjugated anti-GFP antibodies. Free sYFP (3xHA:sYFP) and empty vector (e.v.) were included as controls. The theoretical protein size (in kDa) of each construct is indicated in parentheses. Ponceau S solution staining of RuBisCO was used as a loading control. Three independent experiments were performed with similar results. The results of only one experiment are shown.

**Supplemental Figure 5.**
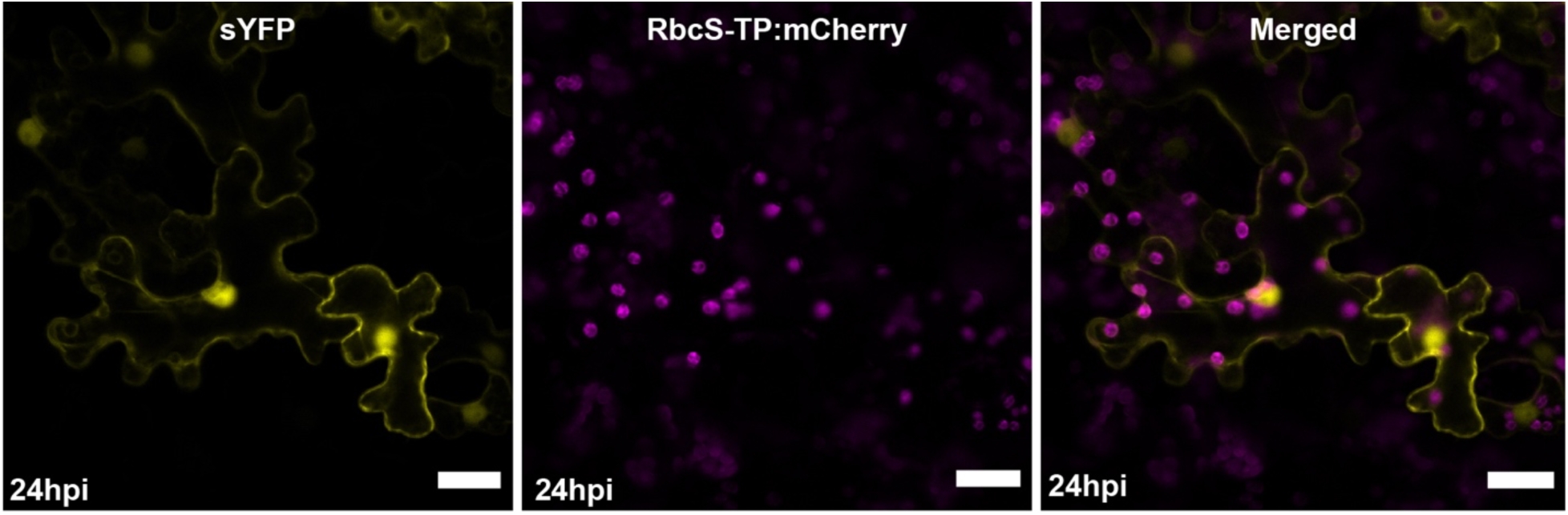
Fluorescence signal from free sYFP is not detected in the stroma of chloroplasts when transiently expressed *N. benthamiana* epidermal cells. Twenty-four hours following agroinfiltration, leaf sections of *N. benthamiana* expressing free sYFP and RbcS-TP:mCherry (plastid marker) were excised and imaged using laser-scanning confocal microscopy. Saturation of the sYFP fluorescent signals revealed no observable protein accumulation in the stroma of chloroplasts. Confocal micrographs shown are single optical sections. The scale bar represents 20 µM.

## Notes

**Funding:** This work was supported by the United States Department of Agriculture, Agriculture Research Service (USDA-ARS) research project 5020-21220-014-00D and an appointment to the Oak Ridge Institute for Science and Education (ORISE). This research was also funded by the Foundation for Food and Agriculture Research New Innovator Award and Indiana Hatch Funds (IND011293) awarded to A.I.P.

### Competing Interest Statement

The authors have declared no competing interest.

## LITERATURE CITED

Ahmed, M.B., Santos, K.C.G.d., Sanchez, I.B., Petre, B., Lorrain, C., Plourde, M.B., Duplessis, S., Desgagne-Penix, I., and Germain, H. 2018. A rust fungal effector binds plant DNA and modulates transcription. Sci. Rep. 8, 14718.

Alfano, J.R. 2009. Roadmap for future research on plant pathogen effectors. Mol. Plant Pathol. 10: 805–813.

Boch, J., and Bonas U. 2010. Xanthomonas AvrBs3 family-type III effectors: discovery and function. Annu. Rev. Phytopathol. 48: 419–436.

Caillaud, M. C., Piquerez, S. J. M., Fabro, G., Steinbrenner, J., Ishaque, N., and Beynon, J., Jones, J.D.G. 2012. Subcellular localization of the Hpa RxLR effector repertoire identifies a tonoplast-associated protein HaRxL17 that confers enhanced plant susceptibility. Plant J. 69: 252–265.

Denne, N.L., Hiles, R.R., Kyrysyuk, O., Iyer-Pascuzzi, A.S., and Mitra, R.M. 2021. *Ralstonia solanacearum* effectors localize to diverse organelles in *Solanum* hosts. Phytopathology 111: 2213–2226.

Earley, K.W., Haag, J.R., Pontes, O., Opper, K., Juehne, T., Song, K., and Pikaard, C.S. 2006. Gateway-compatible vectors for plant functional genomics and proteomics. Plant J. 45: 616–629.

Edgar, R.C. 2004. MUSCLE: multiple sequence alignment with high accuracy and high throughput. Nucleic Acids Res. 32: 1792–1797.

Fabro, G. 2022. Oomycete intracellular effectors: specialised weapons targeting strategic plant processes. New Phytol. 233: 1074–1082.

Felsenstein, J. 1985. Confidence limits on phylogenies: an approach using the bootstrap. Evolution 39: 783–791.

Figueroa, M., Ortiz, D., and Henningsen, E.C. 2021. Tactics of host manipulation by intracellular effectors from plant pathogenic fungi. Curr. Opin. Plant Biol. 62: 102054.

Gaouar, O., Morency, M.J., Letanneur, C., Seguin, A., and Germain, H. 2016. The 124202 candidate effector of *Melampsora larici-populina* interacts with membranes in Nicotiana and Arabidopsis. Can. J. Plant Pathol. 38: 197–208.

Gu, Y., and Innes, R.W. 2011. The KEEP ON GOING protein of Arabidopsis recruits the ENHANCED DISEASE RESISTANCE 1 protein to trans-golgi network/early endosome vesicles. Plant Physiol. 155: 1827–1838.

Guo, B., Wang, H., Yang, B., Jiang, W., Jing, M., Li, H., Xia, Y., Xu, Y., Hu, Q., Wang, F., Yu, F., Wang, Y., Ye, W., Dong, S., Xing, W., and Wang, Y. 2019. *Phytophthora sojae* effector PsAvh240 inhibits host aspartic protease secretion to promote infection. Mol Plant 12: 552–564.

Helm, M., Qi, M., Sarker, S., Yu, H., Whitham, S.A., and Innes, R.W. 2019. Engineering a decoy substrate in soybean to enable recognition of the *Soybean mosaic virus* NIa protease. Mol. Plant Microbe Interact. 32: 760–769.

Helm, M., Singh, R., Goodwin, S.B., Caldwell, D., and Iyer-Pascuzzi, A.S. 2022. Tar Spot of Maize: Current knowledge of genetic interactions and future research prospects to improve disease resistance. Authorea. [Preprint] Available at: https://doi.org/10.22541/au.164642664.47870546/v2

Jones, J., and Dangl, J. 2006. The plant immune system. Nature 444: 323–329.

Kamoun, S. 2007. Groovy times: filamentous pathogen effectors revealed. Curr. Opin. Plant Biol. 10: 358–365.

Kemen, E., Kemen, A. C., Rafiqi, M., Hempel, U., Mendgen, K., Hahn, M., and Voegele, R.T. 2005. Identification of a protein from rust fungi transferred from haustoria into infected plant cells. Mol. Plant Microbe Interact. 18: 1130–1139.

Littlejohn, G.R., Breen, S., Smirnoff, N., and Grant, M. 2021. Chloroplast immunity illuminated. New Phytol. 229: 3088–3107.

Lorrain, C., Petre, B., and Duplessis, S. 2018. Show me the way: rust effector targets in heterologous plant systems. Curr. Opin. Microbiol. 46: 19–25.

Ma, L., Lukasik, E., Gawehns, F., and Takken, F.L.W. 2012. The use of agroinfiltration for transient expression of plant resistance and fungal effector proteins in *Nicotiana benthamiana* leaves. Methods Mol. Biol. 835: 61–74.

Mottaleb, K.A., Loladze, A., Sonder, K., Kruseman, G., and San Vicente, F. 2019. Threats of tar spot complex disease of maize in the United States of America and its global consequences. Mitig. Adapt. Strateg. Glob. Chang. 24: 281–300.

Mueller, D. S., Wise, K. A., Sisson, A. J., Allen, T. W., Bergstrom, G. C., Bissonnette, K. M., Bradley, C.A., Byamukama, E., Chilvers, M.I., Collins, A.A., Esker, P.D., Faske, T.R., Friskop, A.J., Hagan, A.K., Heiniger, R.W., Hollier, C.A., Isakeit, T., Jackson-Ziems, T.A., Jardine, D.J., Kelly, H.M., Kleczewski, N.M., Koehler, A.M., Koenning S.R., Malvick, D.K., Mehl, H.L., Meyer, R.F., Paul, P.A., Peltier, A.J., Price, P.P., Robertson, A.E., Roth, G.W., Sikora, E.J., Smith, D.L., Tande, C.A., Telenko, D.E.P., Tenuta, A.U., Thiessen, L.D., and Wiebold, W.J. 2020. Corn yield loss estimates due to diseases in the United States and Ontario, Canada, from 2016 to 2019. Plant Health Prog. 21: 238–247.

Murphy, F., He, Q., Armstrong, M., Giuliani, L.M., Boevink, P.C., Zhang, W., Tian, Z., Birch, P.R.J., Gilroy, E.M. 2018. The potato MAP3K StVIK is required for the *Phytophthora infestans* RXLR effector Pi17316 to promote disease. Plant Physiol. 177: 398–410.

Nei, M., and Kumar, S. 2000. Molecular evolution and phylogenetics. *Oxford University Press*, *New York*.

Nelson, B.K., Cai, X., and Nebenfuhr, A. 2007. A multicolored set of in vivo organelle markers for co-localization studies in Arabidopsis and other plants. Plant J. 51: 1126–1136.

Petre, B., Lorrain, C., Saunders, D.G.O., Win, J., Sklenar, J., Duplessis, S., and Kamoun, S. 2016. Rust fungal effectors mimic host transit peptides to translocate into chloroplasts. Cell Microbiol. 18: 453–465.

Petre, B., Saunders, D.G.O., Sklenar, J., Lorrain, C., Win, J., Duplessis, S., and Kamoun, S. 2015. Candidate effector proteins of the rust pathogen *Melampsora larici-populina* target diverse plant cell compartments. Mol. Plant Microbe Interact. 28: 689–700.

Petre, B., Win, J., Menke, F.L.H., and Kamoun, S. 2017. Protein-protein interaction assays with effector-GFP fusions in *Nicotiana benthamiana*. Methods Mol. Biol. 1659: 85–98.

Qi, D., DeYoung, B.J., and Innes, R.W. 2012. Structure-function analysis of the coiled-coil and leucine-rich repeat domains of the *RPS5* disease resistance protein. Plant Physiol. 158: 1819–1832.

Redkar, A., Hoser, R., Schilling, L., Zechmann, B., Krzymowska, M., Walbot, V., and Doehlemann, G. 2015. A secreted effector protein of *Ustilago maydis* guides maize leaf cells to form tumors. Plant Cell 27: 1332–1351.

Robin, G.P., Kleemann, J., Neumann, U., Cabre, L., Dallery, J., Lapalu, N., and O’Connell, R.J. 2018. Subcellular localization screening of *Colletotrichum higginsianum* effector candidates identifies fungal proteins targeted to plant peroxisomes, golgi bodies, and microtubules. Front Plant Sci. 9: 562.

Rocco da Silva, C., Check, J., MacCready, J. S., Alakonya, A. E., Beiriger, R. L., Bissonnette, K. M., Collins, A., Cruz, C.D., Esker, P.D., Goodwin, S.B., Malvick, D., Mueller, D.S., Paul, P., Raid, R., Robertson, A.E., Roggenkamp, E., Ross, T.J., Singh, R., Smith, D.L., Tenuta, A.U., Chilvers, M.I., and Telenko, D.E.P. 2021. Recovery plan for tar spot of corn, caused by *Phyllachora maydis*. Plant Health Prog. 22: 596–616.

Rodríguez-Puerto, C., Chakraborty, R., and Singh, R. Rocha-Loyola, P., and Rojas, C.M. 2022. The *Pseudomonas syringae* type III effector HopG1 triggers necrotic cell death that is attenuated by AtNHR2B. Sci. Rep. 12: 5388.

Ruhl, G., Romberg, M.K., Bissonnette, S., Plewa, D., Creswell, T., and Wise, K.A. 2016. First report of tar spot on corn caused by *Phyllachora maydis* in the United States. Plant Dis. 100: 1496.

Saitou, N., and Nei, M. 1987. The Neighbor-Joining Method—a new method for reconstructing phylogenetic trees. Mol. Biol. Evol. 4: 406–425.

Scott, M.S., Troshin, P.V., and Barton, G.J. 2011. NoD: a Nucleolar localization sequence detector for eukaryotic and viral proteins. BMC Bioinform. 12: 317.

Singh, R., Dangol, S., Chen, Y., Choi, J., Cho, Y. S., Lee, J. E., Choi, M., and Jwa, N. 2016. *Magnaporthe oryzae* effector AVR-Pii helps to establish compatibility by inhibition of the rice NADP-malic enzyme resulting in disruption of oxidative burst and host innate immunity. Mol. Cells 39: 426–438.

Schornack, S., Van Damme, M., Bozkurt, T. O., Cano, L. M., Smoker, M., Thines, M., Gaulin, E., Kamoun, S., and Huitema, E. 2010. Ancient class of translocated oomycete effectors targets the host nucleus. PNAS 107: 17421–17426.

Sperschneider, J., Catanzariti, A.M., DeBoer, K., Petre, B., Gardiner, D.M., Singh, K.B., Dodds, P.N., and Taylor, J.M. 2017. LOCALIZER: subcellular localization prediction of both plant and effector proteins in the plant cell. Sci. Rep. 7: 44598.

Sperschneider, J., and Dodds, P.N. 2022. EffectorP 3.0: Prediction of apoplastic and cytoplasmic effectors in fungi and oomycetes. Mol. Plant Microbe Interact. 35: 146–156.

Stam, R., Howden, A., Delgado Cerezo, M., Amaro, T., Motion, G., Pham, J., and Huitema, E. 2013. Characterization of cell death inducing *Phytophthora capsici* CRN effectors suggests diverse activities in the host nucleus. Front. Plant Sci. 4: 387.

Tamura, K., Nei, M., and Kumar, S. 2004. Prospects for inferring very large phylogenies by using the neighbor-joining method. PNAS 101: 11030–11035.

Telenko, D. E. P., Chilvers, M. I., Kleczewski, N. M., Smith, D. L., Byrne, A. M., Devillez, P., Diallo, T., Higgins, D., Joos, D., Kohn, K., Lauer, J., Mueller, B., Singh, M.P., Widdicombe, W.D., and Williams, L.A. 2019. How tar spot of corn impacted hybrid yields during the 2018 Midwest epidemic. Crop Protection Network. doi: 10.31274/cpn-20190729-002.

Telenko, D. E. P., Ross, T. J., Shim, S., Wang, Q., and Singh, R. 2020. Draft genome sequence resource for *Phyllachora maydis*—an obligate pathogen that causes tar spot of corn with recent economic impacts in the United States. Mol. Plant Microbe Interact. 33: 884–887.

Teufel, F., Almagro Armenteros, J.J., Johansen, A.R., Gislason, M.H., Pihl, S.I., Tsirigos, K.D., Winther, O., Brunak, S., von Heijne, G., Nielsen, H. 2022. SignalP 6.0 predicts all five types of signal peptides using protein language models. Nat. Biotechnol. doi.org/10.1038/s41587-021-01156-3

Turnbull, D., Yang, L., Naqvi, S., Breen, S., Welsh, L., Stephens, J., Morris, J., Boevink, P.C., Hedley, P.E., Zhan, J., Birch, P.R., and Gilroy, E.M. 2017. RXLR effector AVR2 up-regulates a brassinosteroid-responsive bHLH transcription factor to suppress immunity. Plant Physiol. 174: 356–369.

Valle-Torres, J., Ross, T. J., Plewa, D., Avellaneda, M. C., Check, J., Chilvers, M. I., Cruz, A.P., Dalla Lana, F., Groves, C., Gongora-Cancul, C., Henriquez-Dole, L., Jamann, T., Kleczewski, N., Lipps, S., Malvick, D., McCoy, A.G., Mueller, D.S., Paul, P.A., Puerto, C., Schloemer, C., Raid, R.N., Roberston, A., Roggenkamp, E.M., Smith, D.L., Telenko, D.E.P., and Cruz, C.D. 2020. Tar Spot: an understudied disease threatening corn production in the Americas. Plant Dis. 104: 2541–2550.

Vargas, W. A., Sanz-Martín, J. M., Rech, G. E, Armijos-Jaramillo, V. D., Rivera, L. P., Echeverria, M. M., Diaz-Minguez, J.M., Thon, M.R., and Sukno, S.A. 2016. A fungal effector with host nuclear localization and DNA-binding properties is required for maize anthracnose development. Mol. Plant Microbe Interact. 29: 83–95.

Wang, Q., Han, C., Ferreira, A. O., Yu, X., Ye, W., Tripathy, S., Kale, S., Gu, B., Sheng, Y., Sui, Y., Wang, X., Zhang, Z., Cheng, B., Dong, S., Shan, W., Zheng, X., Dou, D., Tyler, B., and Wang, Y. 2011. Transcriptional programming and functional interactions within the *Phytophthora sojae* RXLR effector repertoire. Plant Cell 23: 2064−2086.

Wang, X., Zhai, T., Zhang, X., Tang, C., Zhuang, R., Zhao, H., Xu, Q., Cheng, Y., Wang, J., Duplessis, S., Kang, Z., and Wang, X. 2021. Two stripe rust effectors impair wheat resistance by suppressing import of host Fe-S protein into chloroplasts. Plant Physiol. 187: 2530–2543.

Whisson, S. C., Boevink, P. C., Wang, S., and Birch, P. R. J. 2016. The cell biology of late blight disease. Curr. Opin. Microbiol. 34: 127–135.

Win, J., Kamoun, S., and Jones, A.M.E. 2011. Purification of effector-target protein complexes via transient expression in *Nicotiana benthamiana*. Methods Mol. Biol. 712: 181–194.

Xu, Q., Tang, C., Wang, X., Sun, S., Zhao, J., Kang, Z., and Wang, X. 2019. An effector protein of the wheat stripe rust fungus targets chloroplasts and suppresses chloroplast function. Nat. Commun. 10: 5571.

Zipfel, C. 2014. Plant pattern-recognition receptors. Trends Immunol. 35: 345–351.

Zheng, X., McLellan, H., Fraiture, M., Liu, X., Boevink, P. C., Gilroy, E. M., Chen, Y., Kandel, K., Sessa, G., Birch, P.R.J., and Brunner, F. 2014. Functionally redundant RXLR effectors from *Phytophthora infestans* act at different steps to suppress early flg22-triggered immunity. PLoS Pathog. 10, e1004057.

